# Genome-wide mapping of Bicoid/DNA interactions reveals quantitative constraints on transcriptional regulation

**DOI:** 10.64898/2026.05.22.727281

**Authors:** Sadia Siddika Dima, Gregory T. Reeves

## Abstract

During *Drosophila* embryogenesis, Bicoid (Bcd) forms a gradient that provides positional information to regulate target genes along the anteroposterior axis. To understand how the information provided by the Bcd gradient drives gene expression in a concentration-dependent manner, the subpopulations of Bcd participating in gene regulation need to be characterized. Therefore, to understand the mechanism of Bcd-mediated gene regulation, we quantified the absolute concentration of the nuclear subpopulations of Bcd, such as freely diffusing and DNA bound Bcd. These subpopulations have distinct diffusivities and DNA-binding properties and are crucial for gene regulation. This quantification allowed us to construct a global dose/response relationship between the free and DNA-bound concentration of Bcd. Our data show that Bcd/DNA binding is strongly correlated with the free concentration of Bcd, indicated by the dose/response relationship being in the linear regime, despite the barrier presented by nucleosomes. Our data are quantitatively consistent with a Monod-Wyman-Changeux model in which transcription factors passively compete with nucleosomes for DNA binding. We further apply this model to Bcd/DNA interactions at the enhancer/promoter for *hunchback* (*hb*), a Bcd target gene which has a steep posterior boundary, despite being driven primarily by the graded Bcd concentration, a conundrum which has been under scrutiny for decades. We show that, using parameters determined from the global dose/response relationship, a reversible multistate promoter model, in which the promoter activation rate is determined by Bcd binding to the *hb* enhancer, can successfully recapitulate features of *hb* transcriptional dynamics, including the sharp posterior boundary. Therefore, this work sheds light on mechanism of *hb* regulation by Bcd and provides a potential experimental/computational pipeline that bridges the input of global properties of transcription factors to the transcriptional output of specific target genes.

## Introduction

Patterning in multicellular organisms is directed by the graded spatial distribution of morphogens. One of the best-studied morphogens (and the first experimentally-confirmed morphogen) is Bicoid (Bcd), which is a transcription factor (TF) that forms a gradient to pattern the anterior-posterior (AP) axis in the *Drosophila* embryo^1,2^. *bcd* mRNA is deposited maternally in the egg and localized in the anterior pole^1,2^. After fertilization, Bcd protein is translated, and it diffuses outward through the syncytial embryo from its source at the anterior pole to give rise to an exponential anterior-posterior gradient^1,2^. The gradient is believed to regulate target genes, such as the gap genes involved in segment determination in early *Drosophila* development^3^, in a concentration-dependent manner^4^. The Bcd gradient is stable during nuclear cycles 10-14 despite the highly dynamic events during mitosis^5,6^ and precise gene expression patterns are rapidly established in response^7,8^. To explain how a graded morphogen can lead to spatially distinct domains of gene expression, the classical gradient-affinity model was proposed. This model was based on the initial observation that enhancers with weak Bcd binding sites could only drive expression in the anterior regions of the embryo where the Bcd concentration is high, whereas enhancers with strong affinity Bcd binding sites could also drive expression more posteriorly where Bcd concentration is lower^9^. However, this view was soon challenged as regulatory enhancers for more target genes were studied^10,11^. A model has been proposed in which Bcd can activate genes at the anterior pole at high concentrations, while at lower concentrations, Bcd must act in combination with other TFs, such as Hunchback and/or Caudal, to regulate target genes^12,13^. Recent studies of the temporal interpretation of the morphogen also suggest the anterior gap genes have a higher reliance on Bcd, requiring persistent Bcd inputs for activation and maintenance^14^.

How the smooth, graded distribution of Bcd is interpreted into sharp borders of targe gene expression remains unclear. For example, the expression pattern of the target gene *hunchback* (*hb*) is precise with a steep posterior border, thereby neighboring nuclei must be able interpret the small difference in Bcd concentration to result in two distinct responses^15–17^. To address this question, it has been proposed that cooperative binding of Bcd to DNA leads to the sharpness in target gene expression^10,15,16^. The latest advances suggest Bcd forms hubs that facilitate DNA binding by increasing the local concentration, especially in the locations with low Bcd concentration^18,19^. These hubs have also been proposed to act as fast sensors of positional information provided by the concentration gradient^20^. Additionally, slowly moving clusters (non-DNA-bound) have been observed, and are proposed to provide an efficient search strategy for the rapid and precise transcriptional response^21^. The pioneer factor Zelda (Zld) has also been proposed to facilitate the binding of Bcd, particularly at low concentrations of Bcd^19,22^. Interestingly, Bcd itself has been proposed to show pioneer-like activity at high concentrations, thereby modifying chromatin structure at target enhancers^23,24^. This indicates a complex interplay between the Bcd concentration gradient, binding sites with varying affinities, and chromatin structure to regulate gene expression.

To understand the mechanism by which the Bcd concentration gradient conveys positional information to target genes, a quantitative distinction must be made among the various subpopulations of Bcd including, at minimum, the free and DNA-bound populations. Quantitative measurements of the relative and absolute concentrations of the Bcd gradient^5,15,25^, although important to understand the morphogen behavior, are not sufficient to explain its activity. Similarly, although local measurements at the transcriptional hubs using advanced techniques like the lattice light-sheet microscopy brought another important aspect of gene regulation to light^18,19^, a connection remains to be made between nucleus-level Bcd concentration and local, enhancer-specific occupancy. Furthermore, target gene expression appears to be a complex, unknown function of the TF occupancy, which in turn is a complex function of the free concentration. Thus, the dose/response relationships between the free concentration and DNA-bound Bcd, as well as between DNA-bound Bcd and gene expression, remain uncharted.

To address these gaps in knowledge, we used Raster Image Correlation Spectroscopy (RICS)^26–32^, a type of scanning Fluorescence Correlation Spectroscopy, which allowed us to quantify the nucleus-averaged absolute concentration of subpopulations of Bcd with distinct diffusivities, representing populations that are immobile due either to DNA-binding or to formation of clusters. As previously, we extended this methodology to long time course RICS (“tRICS”) acquisitions^31,32^, mapping these biophysical parameters over time and AP location in the blastoderm stage embryo. Our results suggested that the positional information provided by the Bcd gradient is preserved in the subpopulations of Bcd. At the genome-level, the dose/response relationship between free and DNA-bound Bcd indicates cooperative binding of Bcd, which can be explained by the Monod-Wyman-Changeux (MWC) mathematical model of nucleosome-mediated cooperativity between Bcd molecules. In this model, Bcd at high concentration can compete indirectly with nucleosomes, leading to cooperative DNA binding and replacement of nucleosomes.

In order to investigate how the genome-level information is interpreted at a specific target gene locus, we considered the target gene *hb*, which has been visualized by the MS2-MCP approach^8^. We combined a thermodynamic enhancer occupancy model with a multistate promoter model in which the promoter has to overcome kinetic barriers before transcription can initiate. We found that a model in which the promoter transitions reversibly among multiple transcriptionally ON/OFF states could recapitulate the features of transcriptional dynamics, such as the switch-like pattern of the fraction of active nuclei and the transcriptional onset time, using the parameters obtained from the genome-wide dose/response relationship. Thus, the dose/response relationship enhances our understanding of how Bcd-DNA binding is dictated by the morphogen concentration at a nucleus-wide level, and the enhancer/promoter model elucidates the mechanism by which the global information is interpreted at the target gene regulatory region to drive gene expression.

## Results

### Temporal measurements show rapid DNA binding upon mitotic exit

We used tRICS to quantify the absolute nuclear concentration of Bcd at varying AP locations from nc 10 to 14 (Fig. 1A). Within each nuclear cycle interphase, the absolute nuclear concentration of Bcd initially increases, taking ~ 4 minutes upon mitotic exit to reach the peak concentration, after which it begins to decrease (Fig. 1B). The concentration is restored to a similar level in the beginning of the next interphase, despite the changes in number of nuclei and nuclear volume^6^. Thereby the consistent levels of Bcd over the ncs allows it to provide positional information about the nuclei^5^. The decrease in the concentration during the interphase is likely due to the increase of nuclear volume during the time^5^. Nuclei in all the AP locations follow a similar trend of concentration change over time, with the nuclei far from the anterior pole having lower overall concentration than those near the anterior pole.

**Figure 1:**
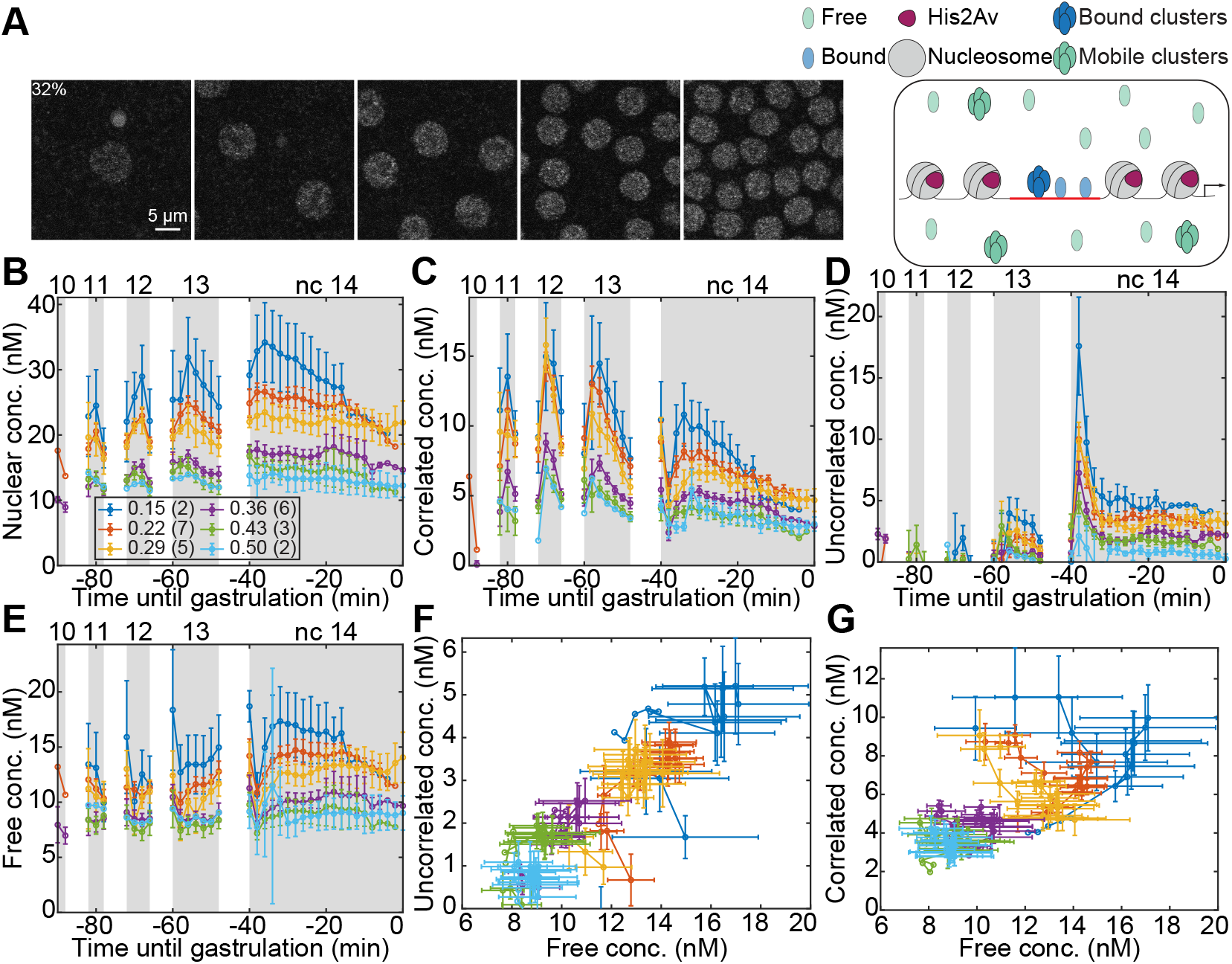
Spatiotemporal dynamics of the populations of Bcd during nc 10 to gastrulation determined using tRICS. (A) Representative images of tRICS acquisitions from nc 10 to 14 (left to right) at the %EL location indicated at the top left. Right panel, illustration of the subpopulation of Bcd in these images characterized using RICS (modified from ^32^, distributed under a Creative Commons Attribution License 4.0 (cc BY)). (B-E) Dynamics of the Bcd populations at different locations with each color denoting a particular location of the region of interest (ROI) with total nuclear concentration (B), concentration of the correlated population (C), concentration of the uncorrelated population (D), and concentration of the free population (E). (F-G) Dose/response relationship between the bound populations and free populations with each color denoting the same locations as in (B-E), for free and uncorrelated populations (F) and free and correlated populations (G). Data points, mean; error bars, SEM. Numbers in parentheses in the legends in (B) indicate number of embryos imaged for each position.

To see how the DNA binding and hub formation of Bcd are related to the free concentration, we quantified the concentration of Bcd that is immobile (or having very low diffusivity). This pool of Bcd is composed of two separate populations: one that correlates with His2Av (referred to as the correlated pool) determined by cross-correlation RICS and the other which does not (referred to as the uncorrelated pool)^31,33^ (Fig. 1A). The concentrations of the correlated and uncorrelated populations are obtained from the product of the total nuclear concentration and the fraction of total Bcd that correlates with His2Av-RFP (Fig. S1A) and the fraction of immobile Bcd that does not correlate with His2Av-RFP (Fig. S1B), respectively. The correlated pool is likely to be the DNA bound pool^34,35^, whereas the uncorrelated pool might be due to the formation of clusters having low diffusivity, as reported previously^21^. We found that, at the start of each nc interphase, the concentration of the correlated population begins to increase, peaking at roughly 6 min after entering interphase, or a ~2 min lag behind the total concentration (Fig. 1C). After reaching a maximum, the concentration of the correlated population starts decreasing, likely due to an increase in nuclear volume having a dilution effect, similar to what is observed in the case of the total nuclear concentration. These observed dynamics demonstrate a prompt, albeit slightly delayed, resumption of transcriptional activity for the rapid and precise establishment of target gene patterns^8^, despite the interruption in the transcriptional activity due to frequent mitoses in the blastoderm-stage embryo. In particular, the brief period at the start of interphase in which the correlated concentration steadily increases may correspond to an overall increase in the chromatin accessibility during interphase, and is consistent with the case of *hb* P2 enhancer accessibility, which has been shown to start increasing 3 min into interphase, reaching a maximum during prophase^36^. The uncorrelated population, which may represent slowly-moving clusters of Bcd, and may also be essential for gene regulation^21^, also demonstrates a similar trend (Fig. 1D).

The free concentration, which acts as the input for DNA-binding and cluster formation, shows a slight increase over the ncs, with highest concentration reached at nc 14 (Fig. 1E). Clear dose/response relationships between the free subpopulation and the two immobile subpopulations are apparent for the data points at different AP locations (see gross trends across data points of different colors in Fig. 1F-G). However, fluctuations in concentration from one nc to the next blur the relationships. This led us to investigate the subpopulations and their resulting dose/response relationships along the AP axis, but within a narrow time window: mid-to-late nc 14, when the subpopulations reach a pseudo-steady state.

### Spatial variation indicates the positional information is preserved in bound populations

To study how the Bcd concentration gradient drives the formation of the correlated and uncorrelated populations, we performed RICS analysis on images acquired at different AP locations during nc 14 (Fig. 2A). Due to the dynamic nature of the subpopulations in early nc 14, we acquired data only between ~10 minutes into nc 14 and gastrulation. During this time window, the dynamics of the subpopulations appear to be at a pseudo-steady state, as the reduction of concentration after reaching a peak value has been shown to be driven by the increase in nuclear size^5^ (Fig. 1B-E). The nuclear concentration shows the well-characterized exponential decay with distance from the anterior pole (Fig. S2A). We found the length constant of the exponential decay curve, λ to be ~0.3, which is slightly higher than the value of 0.2-0.28 reported in fixed embryos by immunofluorescence staining^37,38^. The slight increase in case of our *in vivo* measurements might result from the fact that not all Bcd-GFP is visible in the live embryos due to the time taken for GFP to mature^39^. As such, we applied a correction previously used for this Bcd-GFP construct that accounts for the maturation time of GFP^39^ (see Methods), and the length constant of the total nuclear concentration reduced to 0.23, a value comparable to previous antibody staining. If the subpopulations are key in the Bcd-mediated regulation of the target genes, the positional information of the Bcd gradient must be preserved in these populations. In agreement with this, we found that the concentrations for all the populations follow similar exponential decay along the AP axis (Fig. S2A-G), the maturation correction resulted in slightly lower length constants for each population (Fig. 2B-F).

**Figure 2:**
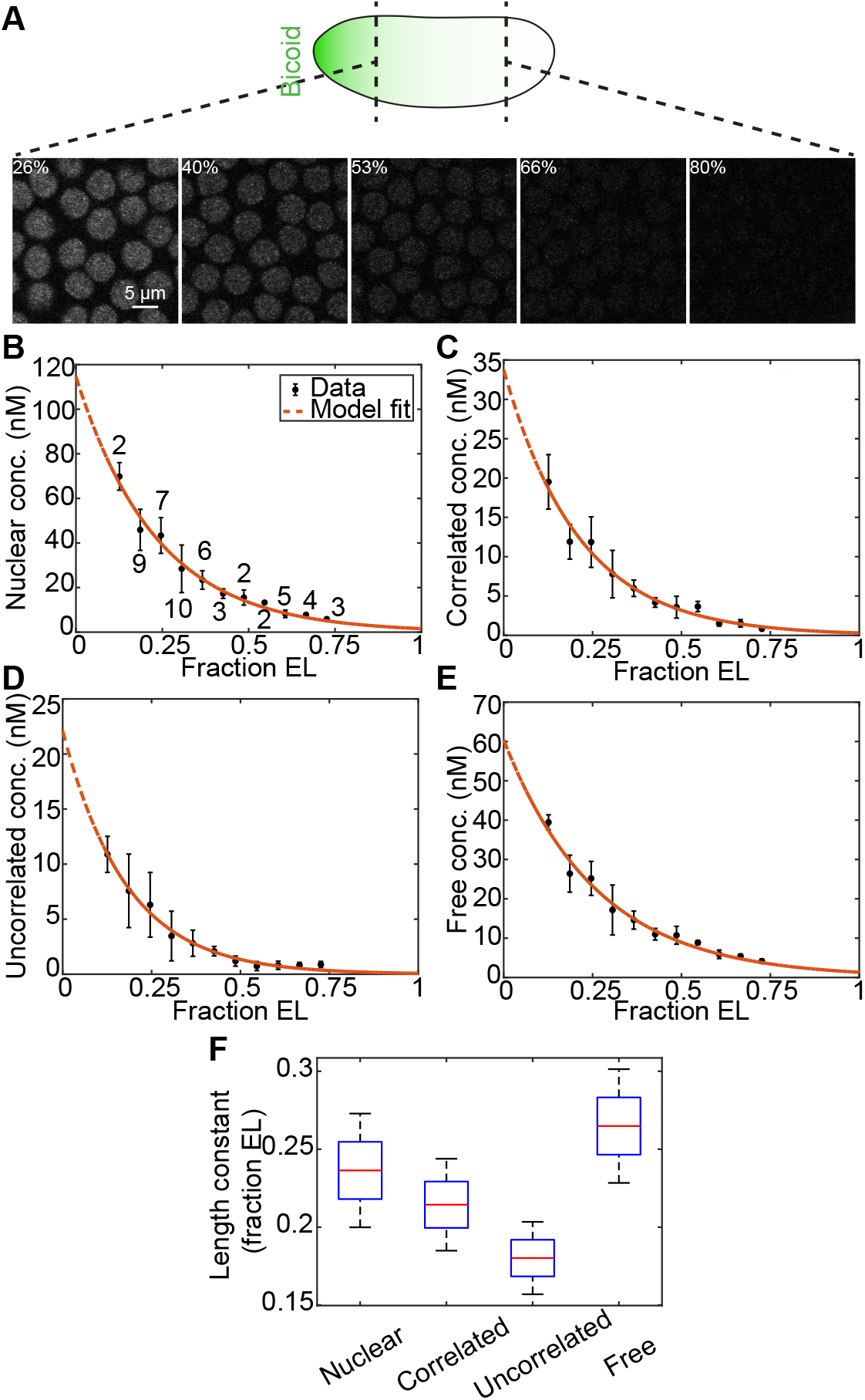
Spatial variation of the populations of Bcd during nc 14 determined using tRICS. (A) Representative images of tRICS acquisitions at different locations along the AP axis. The locations are indicated at the top left of each image and connected to the illustration at top of overall Bcd gradient in the embryo. (B-E) Exponential decay fit to the GFP maturation corrected AP profile of each population including total nuclear concentration (B), concentration of the correlated population (C), concentration of the uncorrelated population (D), concentration of the free population (E). Data points, mean; error bars, SEM. Numbers on the curve in (B) indicate number of embryos imaged for each position. (F) Comparison of the length constant for the exponential decay fit to each population.

We found that the fractions of the correlated and uncorrelated populations show a slightly reducing trend as the distance from the anterior pole increases (Fig. S2E-F), indicating that, despite the presence of Bcd in the more posterior nuclei, a lower fraction of Bcd is binding to DNA in that region of the embryo. As a result, the concentration of the correlated and uncorrelated populations also show approximately exponential decay, similar to the total concentration gradient, but with the shorter length scales of 0.21 and 0.18 respectively (Fig. 2C-D). Correspondingly, we found that the length constant for the free concentration profile is 0.26, which is larger than both the total concentration and the bound populations (Fig. 2F), indicating that the free population does not decline as rapidly along AP axis. This also indicates a pitfall in using the total nuclear concentration and the free concentration interchangeably in thermodynamic equilibrium models.

### Dose/response relationships indicate cooperative binding

The variation of the length constants for the exponentially decaying concentrations of the subpopulations led us to investigate the dose/response relationship between the concentration of the free population and that of the populations with low diffusivities, i.e. the correlated and uncorrelated populations. Since all three populations exhibit exponential-like decay along the AP axis, the dose/response relationships empirically resemble power laws. However, at high concentrations of free Bcd, the two slowly-moving populations should saturate, and as such, power law relationships and exponential decay would not hold. Therefore, even though the free Bcd concentration is not high enough to observe saturation effects, we argue that any empirical or mechanistic model of the slow populations should be a saturable function of free Bcd, such as a Hill or Michaelis-Menten function.

Our data show that the uncorrelated population, likely composed at least in part of the previously-observed clusters of Bcd with low diffusivity^21^, is positively correlated with the free populations. Due to the sigmoidal nature of the data points, we fitted an empirical Hill function to the dose/response relationship (Fig. 3A). The best-fit parameters suggest a Hill coefficient of 2, indicating cooperativity; a dissociation constant of 1.3 µM, indicating low affinity; and a maximum capacity of 20 nM. However, as the identity of this subpopulation, and the mechanism leading to its formation, remain to be investigated, for the rest of the paper we focus on the correlated population.

**Figure 3:**
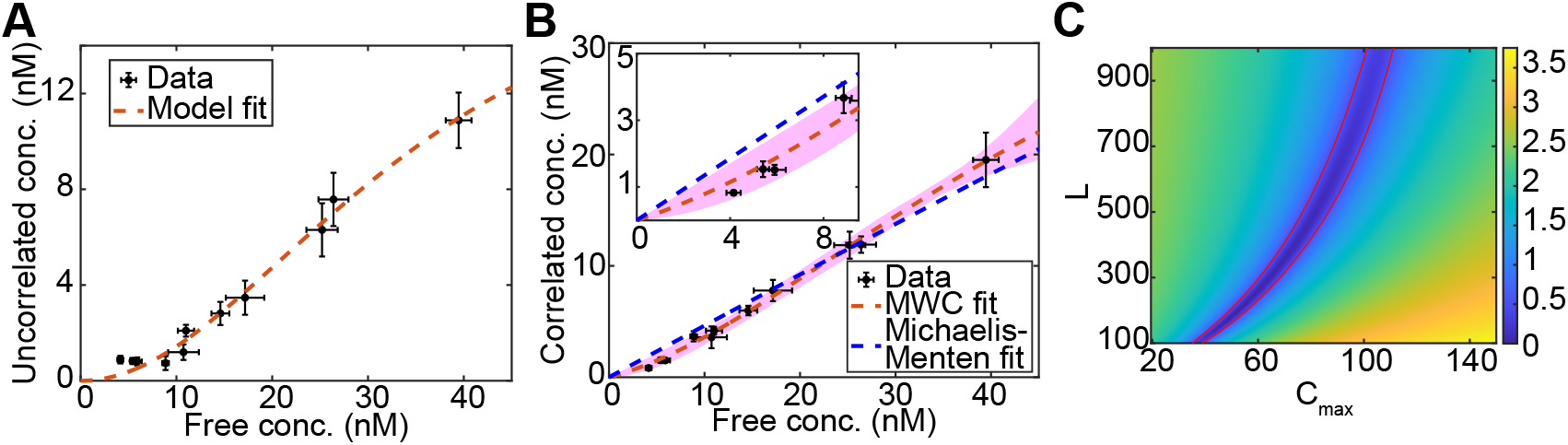
Dose/response relationship between the free and immobile populations at nc 14. (A) Hill function fit to the dose/response between free Bcd and uncorrelated Bcd. Data points, mean; error bars, SEM. (B) Michaelis-Menten and MWC fit to the dose/response between free Bcd and correlated Bcd. The pink shaded region indicates the MWC fit using different parameter sets that fit the data within a set threshold. Inset zooms into the relationship at low free concentration region. Data points, mean; error bars, SEM. (C) Correlation between *L* and *C*_*max*_ when the other parameters are set at the best fit values (*n* = 2, *K*_*N*_ = 320, *K*_*O*_ = 4). Colorbar, log normalized SSE relative to the global minimum.

In contrast to the uncorrelated population, the correlated population likely represents the DNA-bound population of Bcd^34,35^. We found that the dose/response relationship between the free and DNA-bound populations is almost linear, with a positive x-intercept (Fig. 3B). A similar positive correlation has been observed between the nuclear concentration and DNA-bound Bcd clusters^18,20^, which is a component in our correlated population. The positive x-intercept likely indicates that extrapolating back to zero free concentration would require a concave-up portion of the relationship, which could be an indication of cooperative binding of Bcd (Fig. 3B). As such, a simple model of Michaelis-Menten kinetics fails to explain the relationship. Indeed, the cooperative binding of Bcd has been observed and is proposed to be essential for the sharp border of target gene expression in studies focused on specific regulatory regions^7,10,11,40–42^. Our genome-level results suggest that the binding of a Bcd monomer facilitates the successive binding in nearby regions, not only in specific regulatory regions, but also throughout the genome.

### Model of indirect Bcd/nucleosome competition explains cooperativity

Although the cooperative binding of Bcd in regulatory regions is well-known, the mechanism behind it remains unknown. Bcd has been proposed to show pioneer-like activity at high concentrations^23^, which, in conjunction with the cooperative nature of the binding, is congruent with the Monod–Wyman–Changeux (MWC) model of cooperativity^43^. According to this model, which has previously been used for Bcd/DNA binding^24^, a TF can bind its cognate DNA site when the chromatin is either in the open state, or in the nucleosomal state as well (with lower affinity), and the cooperativity of the binding is induced by the competition between nucleosomes and the TF^44^. For TF-DNA binding, the model is as below (see derivation in Methods):

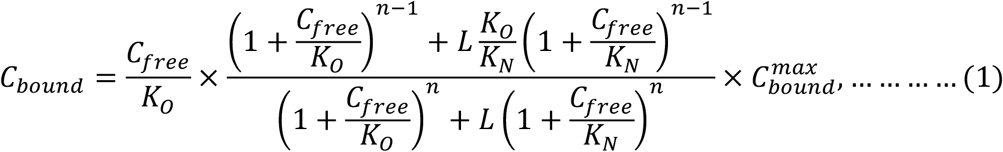

where *C*_*bound*_ is the DNA-bound concentration, *C*_*free*_ is the free concentration, *K*_0_ and *K*_*N*_ are the dissociation constants for binding in the open and nucleosomal states of the DNA respectively, *L* is the equilibrium constant between the two states in absence of bound sites,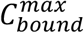 is the maximum capacity for Bcd binding throughout the genome (total concentration of Bcd binding sites), and *n* is the number of binding sites in a given DNA region.

The model input is the concentration of free TF, and the output of DNA-bound TF. However, as these subpopulations are difficult to measure experimentally, especially in live cells, the model has been fit to transcriptional data as a function of total TF nuclear fluorescence, as proxies for the output and input, respectively. For example, previously the model has been fit successfully to the position and steepness of the posterior boundary of *hbP2* expression pattern as a function of Bcd nuclear fluorescent intensity^24^. In contrast, our measurements of the subpopulations of Bcd allowed us to directly use the free and the DNA-bound concentrations, both on a genome-wide level, as the input and output of the model, respectively.

We fit the model to our data using brute-force parameter screening using relevant ranges of the parameters^24,44^ (Table S1) (Fig. S3A-F). The model could explain the observed dose/response behavior, indicating that there is cooperative binding of Bcd at high concentrations (Fig. 3B). The value of *n* is tightly constrained at 2 (Fig. S3A), suggesting that there are 2 Bcd binding sites per cluster on average in the genome.

The value of *K*_*O*_ is also tightly constrained at roughly 2 − 5 *nM* (Fig. S3A-B), which is comparable to the 0.2-5 nM previously found with *in vitro* measurements^11,45,46^ and with Hb expression^47^. *K*_*O*_ being in the low nanomolar range indicates high affinity of Bcd for open (nucleosome-free) DNA sites, and as such, the open DNA sites should be roughly 90% saturated with Bcd at our maximum observed free concentration of ~ 40 nM (Fig. S3G). However, our nucleus-averaged measurements do not approach saturation, remaining instead in the linear regime (Fig. 3B). Therefore, even though the affinity of Bcd for the open sites is high, the fitted model is consistent with maintenance of the linear regime throughout the observed free concentration range of 3 - 40 nM, arguing that saturation is dictated by either the affinity for closed chromatin, *K*_*N*_, or the equilibrium constant, *L*.

In contrast to the tightly-constrained affinity for the open sites, the model maintains consistency with the experimental data for a relatively wide range of affinity for the nucleosome-bound sites, *K*_*N*_. Furthermore, the best-fit parameter range of *K*_*N*_ is 300 – 1500 nM (Fig. S3A-E): large enough values to imply that Bcd does not appreciably bind to closed chromatin at the observed values of free concentration (Fig. S3C-E,H). As such, the indirect competition between Bcd and nucleosomes, parameterized by the equilibrium constant, *L*, is the main driver of the shape (slope of the linear regime and location of the inflection point) of the sigmoidal dose/response curve. Indeed, at high *K*_*N*_, the model can be simplified to neglect the effect of Bcd binding to nucleosome-occluded sites (see Methods). This simplified model also fits our dose/response data well (Fig. S3I) with the parameters falling within the reasonable ranges (Table S1). Furthermore, the location and slope of the inflection point, predicted by the simplified model are very similar to those predicted by the full MWC model (Fig. S3J-K), further indicating the shape of the dose/response curve is controlled primarily by L (see Methods).

In contrast to the other parameters, which are either tightly constrained or largely irrelevant, *L* and 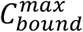 are neither. We found that, upon fitting the full MWC model to our data, the values of 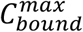 and *L* are highly correlated, and as such, *L* becomes constrained at a fixed value of 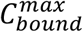 (Fig. 3C). To obtain an estimate of the lower limit of 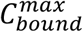, we made use of published Bcd ChIP-seq results (see Methods). Our analysis suggests the concentration of available Bcd binding sites at nc 14 is 29.7 nM, which gives us a lower limit for the capacity. The MWC fits suggest capacity of 30-60 nM and values of L ranging 100-300 fit the experimental data well (Fig. S3A-B).

By performing sensitivity analysis, we also found that the model is less sensitive to changes in L compared to *n, K*_*O*_, and 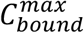, and almost completely insensitive to changes in *K*_*N*_ (Fig. S4A-E). This analysis is consistent with *n, K*_*O*_ being tightly constrained, *K*_*N*_ being unimportant, and *L* having a moderately wide range of values consistent with the data. In the MWC model, it is assumed that, in the absence of TF binding, the open and nucleosomal sites are in equilibrium with equilibrium constant *L*. Therefore, from a reaction equilibrium viewpoint, as Bcd binds the open sites, the equilibrium is shifted such that the closed sites become converted to open sites, indicating there is no direct competition between Bcd and the nucleosomes. Instead, there is indirect competition, with Bcd having the opportunity to bind when the nucleosomes unbind the DNA, as the nucleosomes are dynamic with rapid DNA unwrapping^48^.

The model suggests that passive competition of Bcd with nucleosome leads to cooperative binding at high concentrations. Although there is experimental evidence suggesting Bcd possesses the property to drive chromatin accessibility at high concentrations^23^, we acknowledge that our data cannot rule out the possibility that other mechanisms of cooperativity might also be present. It should also be noted that, our measurements are averaged over the whole nucleus and do not pertain to any particular regulatory region. However, quantification of parameters for genome-wide Bcd-DNA binding led us to investigate whether these can be used to explain the transcriptional dynamics of a target gene driven by a particular regulatory region.

### Bcd occupancy in the enhancer cannot explain hb expression pattern

The step-like activation of the target gene *hb*, driven primarily by the activator Bcd, has been studied for decades^7,17,24,49,50^. The recent advances in the MS2 system^51^ enabled the visualization of nascent transcription, and thereby the elucidation of the transcriptional dynamics of *hb* driven by the hunchback P2 minimal enhancer^7,24,49,50^. The ON/OFF *hb* pattern resembles a Hill function with high Hill coefficient, presumably resulting from cooperative binding^7,24,37,40,41,49,52^. However, an accurate model of *hb* spatial patterning that accounts for mechanistic Bcd/DNA interactions remains elusive.

We consider a thermodynamic enhancer state model for Bcd-mediated regulation of *hb*^49,50,53–56^, which retains the same Bcd/DNA/nucleosome interactions described in the MWC model. In the model, Bcd can bind the open and nucleosomal DNA with affinities *K*_*O*_ and *K*_*N*_, with *K*_*O*_ ≪ *K*_*N*_, and the unbound open and nucleosomal sites are in equilibrium, with equilibrium constant *L*. However, it is important to note that, in the MWC model (Eq. 1), the output is the concentration of Bcd sites occupied by Bcd, whereas, in the enhancer state model, the output is the probability that the enhancer is in an active state, *P*_*active*_.

This probability is given by the ratio of the sum of the active states to the sum of all the states^57^ (Fig. 4A, see Methods for derivation), and as such, one must know the enhancer structure and which states count as “active.” For example, consider a simple case of an enhancer with two Bcd binding sites occluded by a nucleosome in the inaccessible state (Fig. 4A). If the enhancer is considered “active” when any amount of Bcd is bound, the probability of finding the enhancer in an active state is given by

**Figure 4:**
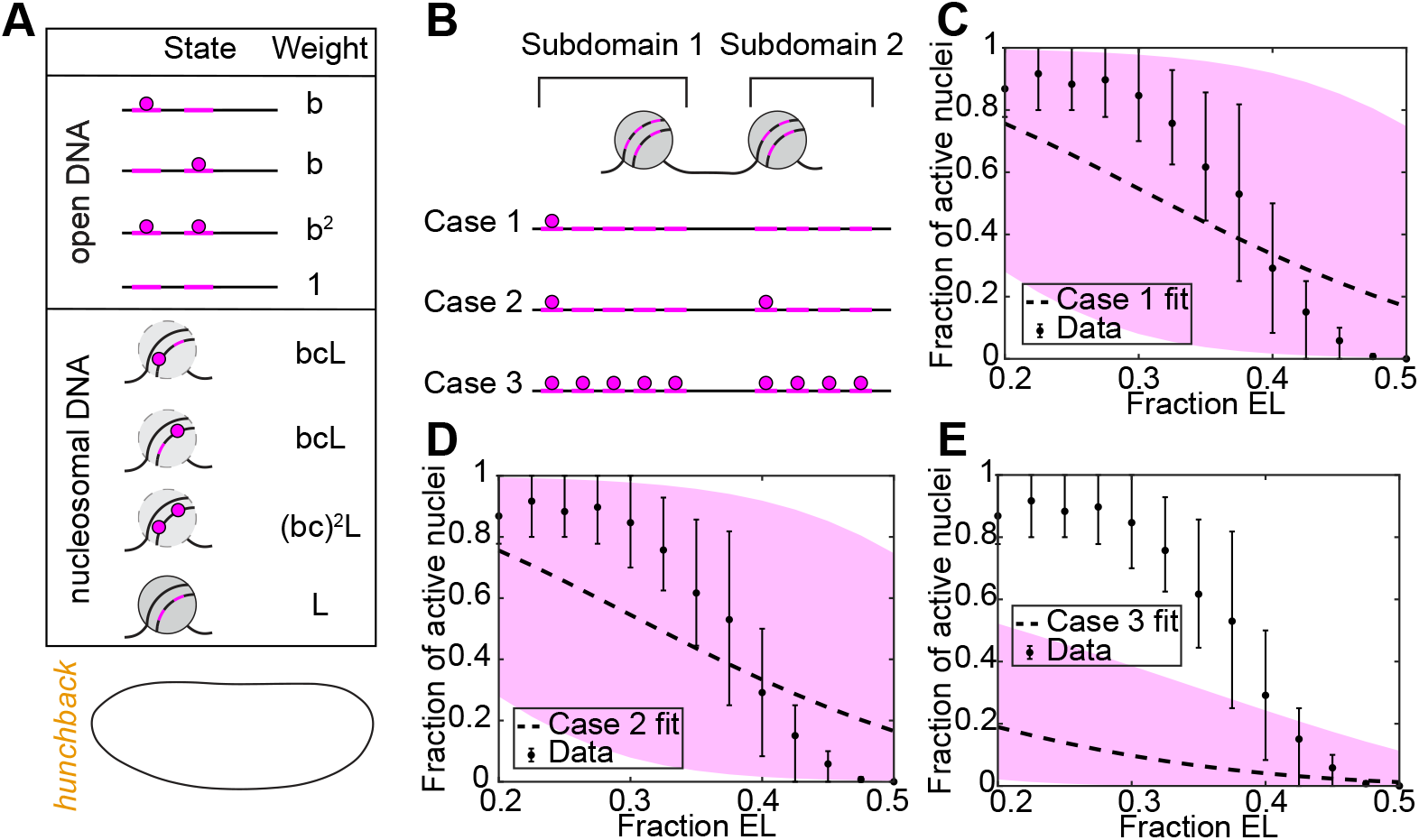
Enhancer state model assuming Bcd occupancy drives gene expression. (A) Illustration of the states of the enhancer and corresponding weights using a simplified case of two Bcd binding sites occluded by a nucleosome in the inactive state. Here, *b* = *C*_*free*_/*K*_*O*_, and *c* = *K*_*O*_/*K*_*N*_. The *hb* expression pattern is illustrated in the bottom panel. (B) Illustration of states corresponding to the minimum number of Bcd molecules bound that drive activation in the three cases of the model for the *hbP2* enhancer. (C-E) The fit to the fraction of active nuclei using the three cases of the model: Case 1 fit (C), Case 2 fit (D), Case 3 fit (E). The dashed curve represents the best fit and the pink shaded region indicates the range of fits using all the parameter sets in the pink shaded region of Fig. 3B. Data points, mean; error bars, inner 68th percentile range.

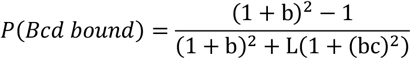

For the *hbP2* enhancer, there are two subdomains: one with 5 Bcd binding sites which can be occluded by a nucleosome, and another with 4 Bcd binding sites which can be occluded by a second nucleosome^24^ (Fig. 4B). As it is unknown which enhancer states are “active,” we considered three cases.

**Case 1**

The enhancer is active as long as any Bicoid is bound

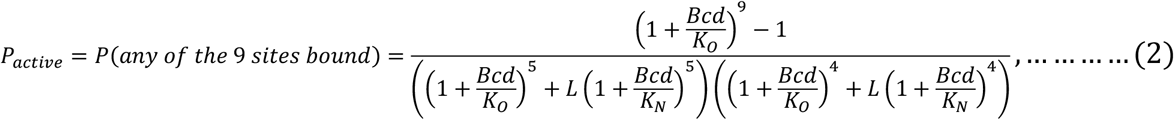

**Case 2**

The enhancer is active as long as at least one Bicoid is bound to each subdomain:

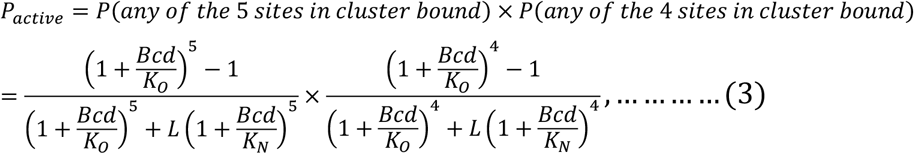

**Case 3**

The enhancer is active only if all the binding sites are bound:

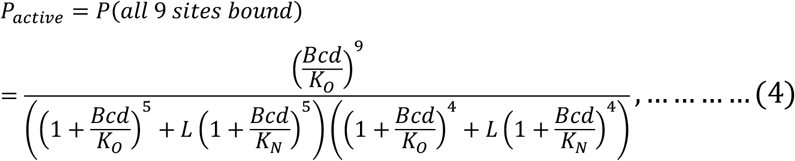

The common assumption for all the cases is that the enhancer is not in the active state if either nucleosome occludes the enhancer. In other words, the enhancer must be in the “open” state for transcription to take place.

Using this model, we investigated whether a simple interpretation in which the probability of transcription is strictly equal to the probability that the enhancer is in the active state could explain the *hb* expression pattern. Using the parameter sets determined from the genome-wide dose/response relationship of Bcd-DNA binding, this model yielded a graded response, and thereby could not recapitulate the *hb* expression pattern such as the position and the step-like pattern of the boundary (Fig. 4C-E). Furthermore, even when we did not constrain the model solely to parameter sets consistent with our genome-wide data, model still could not recapitulate the step-like pattern using any parameter values within reasonable limits (Table S1) (Fig. S5A-C). These results suggest that there may be intermediate steps between TF binding and gene expression, and that TF occupancy in the enhancer is not one-to-one with gene expression (reviewed in ^58^). Given the failure of the enhancer occupancy model to explain the fraction of active nuclei, we considered a multistate promoter model^24,49,59^ that uses the parameters determined from the global dose/response map of Bcd-DNA binding to explain the transcriptional dynamics driven by *hbP2*.

### A reversible promoter model, driven by the enhancer states, can explain the transcriptional dynamics at the target gene locus

In order to explain transcriptional regulation by TF dynamics, recent studies have proposed that promoters overcome kinetic barriers that correspond to one or more inactive states before becoming capable of transcriptional activity^59^ (Fig. 5A). In light of this theory, we consider that the *hb* promoter can transition reversibly among N states, including N-1 OFF states (with probabilities P_1_, P_2_, …, P_N-1_) and one ON state (with probability P_N_). Although the exact number of states that the promoter can be in is unknown, and varying the number of states will slightly change the free parameter values, based on current literature the assumption of three OFF states is reasonable^59^. We assume the rate constant of the forward reactions, *k*_*on*_, is proportional to *P*_*active*_, the probability of finding the enhancer in an active state: *k*_*on*_ = *k*_*on*,0_ *P*_*active*_, where *k*_*on*.0_ is the maximal forward rate constant (Eq. 2-4). We also assume the rate of the reverse reaction (towards the OFF states), *k*_*off*_, is constant (see Methods)^24,49,59^. As above, for the enhancer state model, we make use of the parameters determined using our MWC model fit to the genome-wide dose/response relationship for Bcd-DNA binding (L, *K*_*O*_, and *K*_*N*_). We used this model, informed by our genome-wide data, to investigate two features of *hb* transcriptional dynamics: Fraction of active nuclei and Transcriptional onset time.

**Figure 5:**
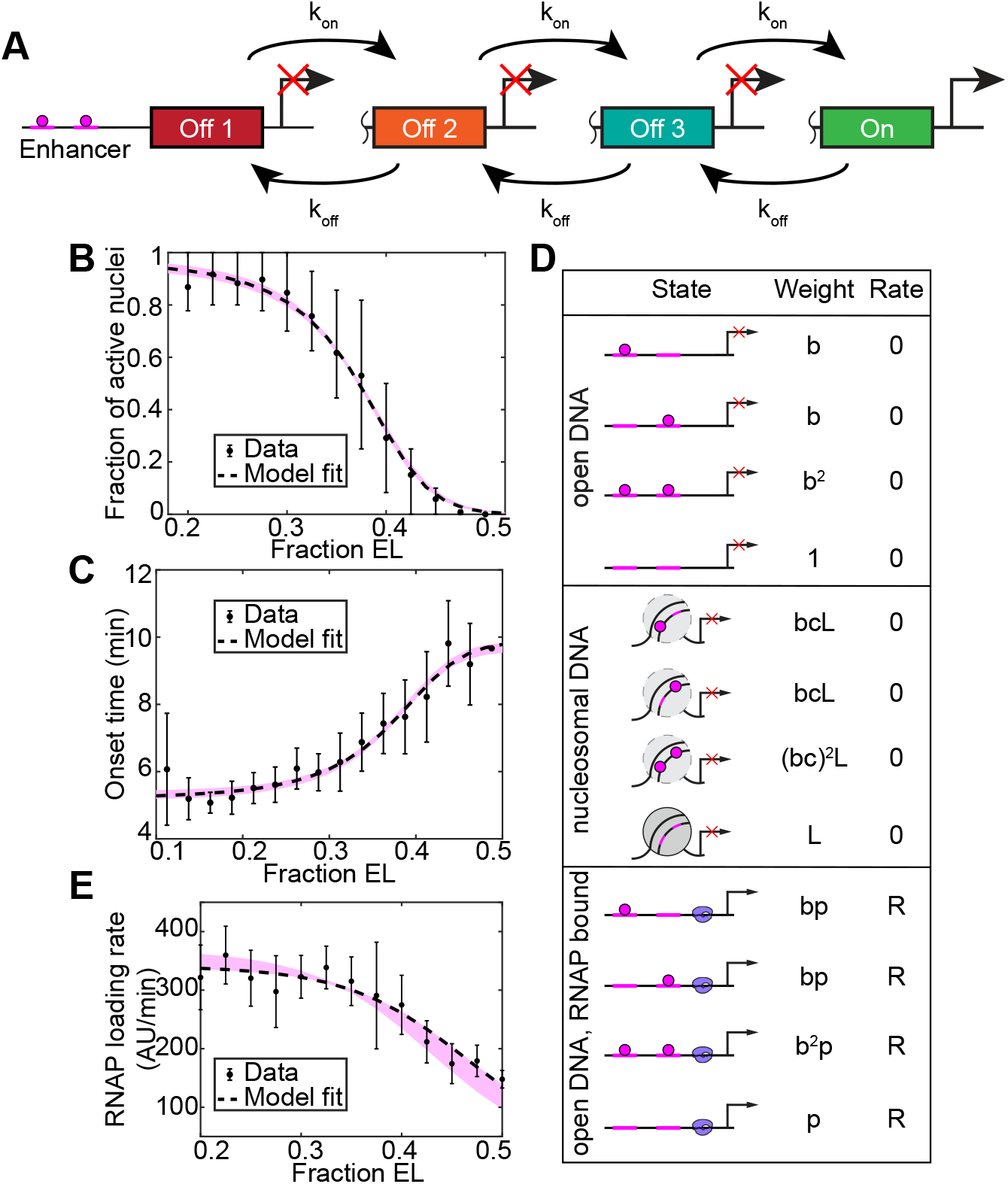
Promoter state model assuming the promoter reversibly transitions between multiple ON/OFF states to drive gene expression. (A) Illustration of the model with three OFF states and one ON state. (B) Model fit to the fraction of active nuclei using Case 1. Dashed curve represents the best fit and the pink shaded region represents the fits using parameter sets in Fig. 3B that fit the fraction of active nuclei within a threshold. (C) Model fit to the transcriptional onset time. Dashed curve represents the best fit and the pink shaded region the fits using parameter sets in Fig. 3B. In (B,C), data points, mean; error bars, inner 68th percentile range. (D) States of the simplified model in Fig. 4A when extended to include the states of RNAP binding assuming the RNAP loading rate is proportional to the probability of finding the promoter in RNAP bound state. (E) Model fit to the RNAP loading rate as reported by Eck et al.^49^. Dashed curve represents the best fit and the pink shaded region the fits using parameter sets in (B). Data points, mean; error bars, SEM.

First, we fit this combined enhancer/promoter state model, at steady state, to the fraction of active nuclei at nc 13, obtained from imaging of embryos with *hbP2-MS2/MCP-GFP* reported by Degen et al.^24^. As the model was fit at steady state, the only remaining parameter, not previously constrained by the MWC model fit to our genome-wide data, is 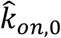, defined as *k*_*on*,0_/*k*_*off*_. We found that this enhancer/promoter state model could tightly recapitulate the fraction of active nuclei (Fig. 5B, Fig. S5D), with the best-fit value of 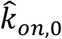 as ~ 35, indicating the transitions towards the ON state are at most 35 fold faster than the transitions away from the ON state. Thus, the model indicates a possible mechanism behind the establishment of the steep ON/OFF pattern of *hb* expression driven by the shallow Bcd gradient.

In these results, we have assumed 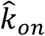 to be proportional to the probability of finding any number of Bcd proteins bound to the enhancer (**Case 1; Eq. 2**). However, since the exact phenomenology that results in activation is unknown, we also investigated whether Cases 2 or 3 of the model are consistent with the data. We found that all three Cases could explain the fraction of active nuclei (Fig. 5B, S5E-F), with slightly different values of the parameter sets (Fig. S5G). Thus, our results indicate that the exact state of the enhancer that drives activation is not as important as the ability of the promoter state model to transform a graded enhancer state model into a steep, switch-like posterior boundary of *hb*. It has been shown, however, that a synthetic reporter with nine Bcd binding sites could recapitulate the spatial features of the expression pattern driven by *hbP2*, whereas reporter with fewer binding sites failed to do so^7^, perhaps suggesting a state of the enhancer in which more Bcd sites are bound might be more favorable to drive activation.

Next, we considered the transcriptional onset time, which is defined as the time at which transcription starts after anaphase, obtained from a previously published database in which embryos with *hbP2-MS2/MCP-GFP* were imaged^24^. As these data are dynamic, it could not be assumed that the model is at steady-state, making the system time scale, *τ* ≡ 1/*k*_*off*_, a free parameter. Using the best-fit value of 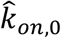 determined from the fraction of active nuclei and the genome-wide parameters *L, K*_*N*_, *K*_*O*_, we fit the enhancer/promoter model to these onset time data (see Methods).

Incorporating a mitotic repression window, defined as the time when the transcriptional machinery remains inactive after mitosis^51^, we found this model was able to recapitulate the onset time, with a best-fit *τ* of ~ 1.6 min and a mitotic repression window of 5 min (Fig. 5C). Together, the values of *τ* and 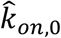 suggest that *k*_*off*_~0.6 min^-1^ and *k*_*on*,0_~21 min^-1^ (Fig. S5G).

Gene expression is a complex function of TF occupancy in the regulatory regions. Hence, the genome-wide MWC model cannot be directly applied to the enhancer since the model input is the TF concentration and the output is TF-DNA binding, not gene expression^44^. Moreover, simpler models assuming that the fraction of active nuclei is proportional to the probabilities of Bcd bound to the regulatory region cannot recapitulate the steplike pattern of fraction active nuclei. Our approach suggests that a shallow-sloped, saturating enhancer state model (Fig. 4B, 5A) is acted upon by a multi-state promoter model to give steplike pattern of *hb*.

### Effect of Zld on hb expression

The pioneer factor Zld acts as a master regulator of zygotic genome activation in *Drosophila*, facilitating the binding of other TFs by establishing and maintaining chromatin accessibility during zygotic genome activation^60,61^. Loss of Zld function has been found to result in reduced Bcd binding in the Bcd-dependent regulatory regions, thereby altering the target gene expression patterns^22,23^. For example, the *hb* posterior boundary shifts to the anterior in *zld* mutant embryos, indicating Zld acts to lower the Bcd concentration threshold for *hb* activation^7,24,49^. In our model, the effect of Zld is incorporated into the equilibrium constant L^24,49^.

If there are *n*_1_ and *n*_2_ Zld binding sites in the subdomains, the probability of Bcd binding in the wildtype embryo as found in Eq. 2 is given by.

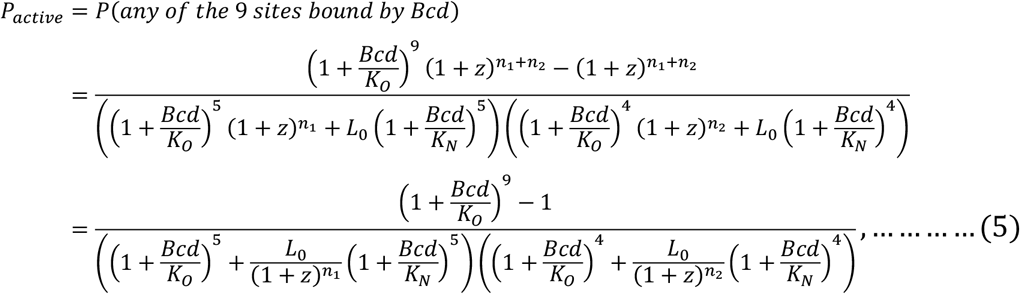

where *z* = [*Zld*]/*K*_*z*_, [*Zld*] is the concentration of free Zld and *K*_*z*_ is the affinity of Zld for DNA. Comparing Eq. (2) and (5), the effective equilibrium constant, *L*, used in the previous sections takes the effect of Zld into account following this relationship: 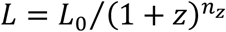, where *L*_0_ is the equilibrium constant in the absence of Zld and *n*_*z*_ is the number of Zld binding sites (see Methods).

For embryos with reduced levels of Zld, such as mutant or RNAi knock-down embryos, the value of the equilibrium constant will be different from that in the wildtype embryos. The probability of finding a Bcd bound state in the embryos with reduced levels of Zld is given by:

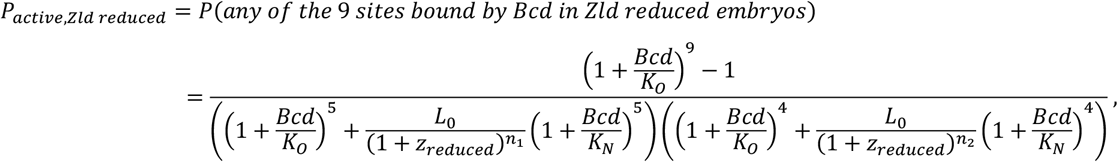

where *z*_*reduced*_= [*Zld*]_*reduced*_/*K*_*z*_ and [*Zld*]_*reduced*_ is the concentration of free Zld in the reduced Zld embryos. The equilibrium constant in the reduced embryos is given by 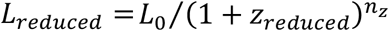. This indicates *L* < *L*_*reduced*_; *since z*_*reduced*_< *z*, which consequently results in an anterior shift of the posterior boundary of *hb* in a reduced Zld embryo^7,24,49^.

### RNAP loading rate can be deduced using the occupancy hypothesis

To use our enhancer state model to predict the RNAP loading rate, which is a key parameter in quantifying gene regulation and can be measured from MS2 data^49–51^, we augment Eq. 2-4 with an additional state: the probability of RNAP binding at the promoter (Fig. 5D, see Methods for derivation)^62,63^. For the ease of visualization, similar to Fig. 4A, we again use the simplified case of an enhancer with two Bcd binding sites occluded by a nucleosome in the inaccessible state to demonstrate the different states that the system can be present in (Fig. 5D). We used these states in the thermodynamic model to calculate promoter occupancy. The enhancer can be either in the accessible or in the inaccessible state. Only the RNAP bound cases can initiate transcription with a rate R.

RNAP binding at the promoter, and thereby transcription, takes place only in the open state, in absence of both the nucleosomes. The probability of RNAP binding to the promoter, *p*_*bound*_, is given by the sum of all the RNAP bound states divided by all the possible states of the system. Therefore, for *hbP2*

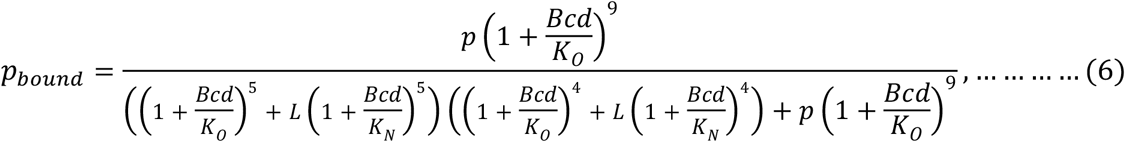

Here, *p* = [*RNAP*]/*K*_*P*_, [RNAP] being the concentration of free polymerase and *K*_*P*_ being the dissociation constant of RNAP.

Then according to the occupancy hypothesis, the rate of mRNA production is given by:

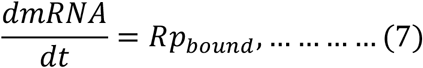

Where R is the rate of transcription when the promoter is bound by the RNAP^49,50^. The slope of the initial rise in RNAP molecules after transcription initiation, as observed from an MS2 trace over time, gives the initial rate of mRNA production or initial RNAP loading rate^49,50^. We fit Eq. 6-7 to a previously-published data set in which the RNAP loading rate was measured^49^. We use the parameter sets (*L, K*_*N*_, *and K*_*O*_) that provide good fit to both the global dose/response and the fraction active nuclei and thus, the normalized RNAP concentration, *p*, was the only adjustable parameter (see Methods). According to our fit, we found that the occupancy hypothesis could explain the graded nature of the RNAP loading rate behavior (Fig. 5E). As such, if indeed the RNAP loading rate is a function of the occupancy, our enhancer state model, pre-constrained by our MWC model fit, is consistent with the previously-published data.

## Discussion

The binding of a TF to the DNA is dependent on a number of factors, including its free concentration, chromatin accessibility, cooperativity, clustering, and interaction with other proteins^58,64–66^. For example, in the early *Drosophila* embryo, DNA-binding properties of many developmental TFs are facilitated by the TF Zld, which has the properties of a pioneer factor, such as the ability to bind nucleosomal DNA, thereby increasing chromatin accessibility^67–71^. In the context of Bcd-mediated patterning, removal of Zld motifs has been found to result in reduced binding of Bcd in the Bcd-dependent enhancers, suggesting that Zld enhances the binding of Bcd, likely through its effect on nucleosomes^72^. As such, recent models of how the bulk Bcd morphogen gradient dictates Bcd/DNA binding along the AP axis have included nucleosome/DNA binding^23,24^. However, measurements of TF/DNA binding, critical to test these models, have been lacking.

To address this issue, in this study we used tRICS to characterize the subpopulations of Bcd composing the bulk concentration. The concentrations of the free population, the correlated population representing the DNA bound monomers and hubs, and the uncorrelated population, all showed exponential decay along the AP axis, similar to the well-studied profile of the bulk Bcd gradient. This indicates the preservation of the positional information provided by the gradient in these populations to a varying extent.

We found that the exponentially decaying concentration profiles resulted in an almost linear dose/response relationship between the free and correlated concentration in the range of concentrations observed in the embryo (Fig. 3B). This suggests that, at a genome-wide level, DNA binding is strongly correlated with the concentration of free Bcd, unlike with the case of the NF-κB transcription factor Dorsal, which also acts during the blastoderm stage^32^. We also found that the binding is suppressed at low concentrations, as apparent from the positive x intercept of the dose/response relationship and the difference between the blue and orange curves at low free concentration (Fig. 3B), indicative of the cooperative binding of Bcd^44^. Indeed, it has been shown that, at high concentration, Bcd can establish accessible states at a subset of its targets that are sensitive to Bcd concentrations and are expressed mainly in the anterior nuclei^23,24^. Although the genome-wide chromatin accessibility is similar in the anterior and posterior halves of the embryo, it has also been found that the anterior enhancers regulated by Bcd are more accessible in the nuclei where they are active^73^, suggesting a role of Bcd in establishing accessibility. This might be explained by an allosteric regulation that induces cooperative TF binding, leading to passive nucleosome eviction^44^.

The cooperative binding of Bcd indicated by our data and also proposed previously^7,10,11,40–42^, as well as the presumed pioneering role of Bcd, led us to use the MWC model of nucleosome-mediated cooperativity^43,44^. The MWC model has been used to infer the relationship between easily accessible measurements, such as the total fluorescence intensity and MS2 transcriptional data^24,49^. Inspired by the success of this model for TF/DNA/nucleosome interactions, we used our detailed quantification of the free and DNA-bound populations, which directly represent the input and output of the MWC model, respectively, to accurately fit the model and estimate parameters associated with Bcd/DNA binding. In turn, analysis of our model fits allowed us to also gain mechanistic insights into the cooperative binding of Bcd. We were able to demonstrate a high affinity of Bcd for open DNA and that it can bind nucleosomal DNA by a principle of passive nucleosome replacement^44^. Furthermore, this model interpretation of passive competition with nucleosomes solves a possible paradox of TFs with high affinities: how a TF can simultaneously have high affinity for its cognate binding sites while also exhibiting differential binding, without saturation, along a wide dynamic range of free concentration. In the MWC model, only a fraction of Bcd binding sites are available for binding, and as Bcd concentrations get higher, the passive competition opens up new opportunities for binding.

The composition of the uncorrelated population remains unknown; however it may include the previously observed slowly-moving clusters^21^, which would be similar to our recent observations of the uncorrelated population of Dorsal^32^. However, unlike Dorsal, the uncorrelated population of Bcd shows a strong dependence on the free concentration (Figs. 1F,3A), and as such, there does not appear to be a threshold in the free concentration for the formation of these clusters. A threshold concentration being a key feature of liquid-liquid phase separation^74,75^, this suggests the Bcd clusters are unlikely to be phase separated. Thus, the mechanism of formation of the slowly moving clusters, and any other component of the uncorrelated population, remains to be investigated.

Our results show that the positional information provided by the morphogen gradient is maintained in the DNA binding and cluster formation. Thus, although the target gene expression might be influenced by inputs from other AP patterning factors^23^, the formation of subpopulations of Bcd is driven in a concentration-dependent manner. The dose/response relationship between the free concentration and the other subpopulations essential for transcriptional regulation indicate that mainly the total gradient dictates the formation of these populations, despite the involvement of numerous other factors in eukaryotic gene regulation. The model suggests that the indirect competition of Bcd with the nucleosome can result in the cooperativity among the noninteracting Bcd molecules leading to cooperative binding.

The Bcd-*hb* system has been used as a model to study gene regulation by morphogens for decades. As such, the structure of the the *hbP2* enhancer is well known, affording us with the opportunity to computationally model enhancer states using the parameters estimated from our genome-wide data. While our enhancer state model is built on the same biochemical interactions as the MWC model, the output is different: instead of the concentration of Bcd-bound sites, the output is the probability of the enhancer being in an active state. Furthermore, *hb* transcriptional dynamics driven by the *hbP2* enhancer have been extensively studied using the MS2/MCP system^7,24,49,50,76^, allowing us to test our enhancer state model against previously reported data.

Our first test was to determine if our measured data and thermodynamic enhancer state model was consistent with the sharp step-like posterior boundary of *hb*. Several mechanisms have been proposed to explain how the graded concentration of the activator Bcd along the AP axis of the early *Drosophila* embryo could produce such a sharp response. However, under the assumption that gene expression is directly proportional to the enhancer occupancy by Bcd, the model failed, reinforcing the proposition that gene expression is a complex, non-linear function of the occupancy^44^. In similar situations, promoter state models have been previously used in attempts to model this non-linear function^24,49,59^. These models suggest that the promoter overcomes multiple kinetic barriers by transitioning among one or more OFF states and one ON state, the latter of which is competent to be transcriptionally active^59^. As such, we combined our enhancer state model, informed by our genome-wide measurements, with a reversible multistate promoter model to explain features of gene regulation. While promoter state models have successfully explained some aspects of transcription^24,49,59^, our combined enhancer/promoter state model, informed by our genome-wide data, tightly recapitulates the switch-like *hb* pattern, beginning from a graded Bcd distribution. Our model was also consistent with the previously reported *hb* transcriptional onset time^24,49,50^. Furthermore, our model suggests that the pioneer factor Zld acts to lower the equilibrium constant between unbound open and nucleosomal DNA, consistent with the anterior shift of the posterior boundary of *hb* in *zld* mutant embryos^24^.

In summary, our *in vivo* measurements allowed us to demonstrate how Bcd-DNA binding is driven by the Bcd concentration, revealing how positional information is conveyed at a genome-wide level. We found that gene expression is not directly proportional to the Bcd occupancy at the enhancer. However, using the parameters obtained from the genome-wide dose/response map, we showed that a model in which the promoter has to reversibly overcome multiple kinetic barriers to reach a transcriptionally active state could recapitulate features of transcriptional dynamics of target gene. Our efforts shed light on the mechanism of gene regulation by a morphogen gradient and proposes a pipeline that can be used to understand gene expression using the knowledge of nucleus-averaged TF properties at a genome-wide level.

## Methods

### Fly stocks

The fly stocks used are: *P{w[+mC]=EGFP-bcd}1, y[1] w[1118]; Diap1[th-1] st[1] kni[ri-1] bcd[6] rn[roe-1] p[p]* (Bloomington Drosophila Stock Center (BDSC) #29018). The flies were crossed *w[*]; P{w[+mC]=His2Av-mRFP1}II*.*2* (Bloomington Drosophila Stock Center (BDSC) #23651). Embryo were collected from females of genotype *P{w[+mC]=EGFP-bcd}1, y[1] w[1118]/w[*]; P{w[+mC]=His2Av-mRFP1}II*.*2/+; Diap1[th-1] st[1] kni[ri-1] bcd[6] rn[roe-1] p[p]/TM3*. Flies were raised on a standard cornmeal-molasses-yeast medium at 25°C. Population cages were started with the selected females with males and kept at room temperature. Grape juice plates streaked with yeast paste was placed on the bottom of the cages for oviposition.

### Live imaging

Embryos were collected from the grape juice plates after the plate was kept for an hour in the cage. Embryos were washed into a mesh basket using deionized water and then dechorionated in bleach for 30 s. The embryos were then washed again with deionized water to remove residual bleach. The embryos were mounted in 1% solution of low melt agarose (IBI Scientific, IB70051) in deionized water on a glass bottom Petri dish (MatTek, P35G-1.5-20-C). After the agarose solidified, it was covered with deionized water to prevent from drying.

### Confocal imaging

Confocal imaging was done using Zeiss LSM 900 confocal laser scanning microscope (ZEISS Microscopy, Germany). For observing the temporal variations, acquisitions were started when the embryos reached nc 10 and continued until gastrulation as indicated by nuclear morphology. A C-Apochromat 40x/1.2 water immersion Korr objective, a 488 nm laser for the eGFP, a 561 nm laser for RFP, and GaAsP-PMT detectors were used. A pixel dwell time of 2.06 μs, a frame time of 5.06 s, and a resolution of 1024×1024 pixel were used. This corresponded to a line time of ~5 ms and a pixel size of 31.95 nm. The region of interest was selected at varying distances from the anterior pole for different embryos to get a range of AP positions. After the acquisition, a z-stack of the whole embryo was taken to measure the respective embryo lengths and distance of the ROI from the anterior pole to get the fraction EL position for each acquisition. For observing spatial variations, additional acquisitions were performed at varying distances from the anterior pole. The acquisitions were started after the embryos reached nc 14 to minimize the effect of photobleaching.

### RICS analysis

We performed RICS analysis according to previously-published protocols^29,31,32^. In brief, two dimensional (2D) RICS ACFs were built from data, then fit to a theoretical 2D ACF in a two-step process. First, the fast direction cut of the theoretical 2D ACF, which is Gaussian-shaped, was fit to the fast direction cut of the data-derived 2D ACF, yielding an ACF amplitude at each time point and hence, the average total concentration. Data from shifts of zero and one pixels were avoided due to shot noise and unwanted correlations in the PMT, respectively. Next, with the amplitude held fixed, we fit a two-component theoretical ACF to the entire data-derived 2D ACF, which yielded the fraction of Bcd that is immobile/slowly-moving.

To determine the fraction bound to DNA, we fit a model of the CCF to the CCF data. This yielded the CCF amplitude, which was normalized by the amplitude of the ACF derived from the H2Av-RFP channel. This ratio is proportional to the correlated fraction, with a correction for the differing sizes of the PSFs of the green and red channels.

### Determination of length constant

We fitted an exponential decay model to the concentration (bulk, correlated, uncorrelated, and free) profiles along the embryo length. For each population:

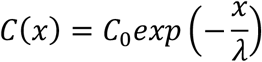

Where, *C*(*x*) is the concentration of the population at a distance of x from the anterior pole, *λ* is the length constant, and *C*_0_ is the maximum concentration at the anterior pole. The values *λ* were determined by brute-force parameter screening and the corresponding values of *C*_0_ were determined analytically.

The time taken for GFP to mature alters the shape of the gradient, resulting in a higher apparent length constant. To take this effect into account, we corrected the observed concentrations:

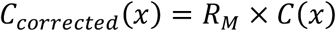

Where *R*_*M*_ is the maturation correction factor for live Bcd-GFP images as obtained by Liu et al.^39^ In this case also, brute-force parameter screening was used to determine the free parameter *λ* and *C*_0_ was determined analytically.

### Empirical model for the dose/response relationship

To explain the dose/response relationships between the free and uncorrelated populations, as well as between free and correlated populations, we considered the Hill function of the form:

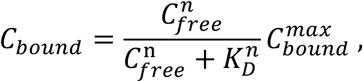

where *C*_*free*_is the free Bcd concentration, *C*_*bound*_ is the correlated or uncorrelated concentration, 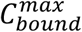 is the corresponding maximum bound Bcd concentration (corresponding to the total concentration of Bcd binding sites in the genome in the correlated case), n is the Hill coefficient representing cooperativity, and *K*_*D*_ is the dissociation constant for Bcd-DNA binding. At *n* = 1, the equation reduces to the Michaelis-Menten kinetics. We used brute-force parameter screening to determine the free parameter *K*_*D*_ in case of Michaelis-Menten and *K*_*D*_ and *n* in case of Hill function. The capacity 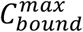 was determined analytically in both the cases.

### MWC model

Derivation (nucleosome-mediated cooperativity)

There are n binding sites in each enhancer, the enhancers can be either in open (O) or nucleosomal (N) state.

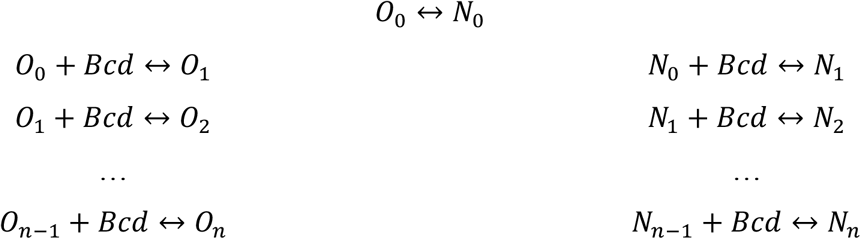

*K*_*O*_ is equilibrium constant for the open state and *K*_*N*_ is that for the nucleosomal state. *C*_*free*_is the concentration of free Bcd.

Now, in the absence of Bcd the two states (open and nucleosomal) are in equilibrium and *L* is the equilibrium constant. Therefore, 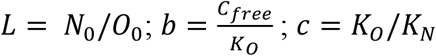

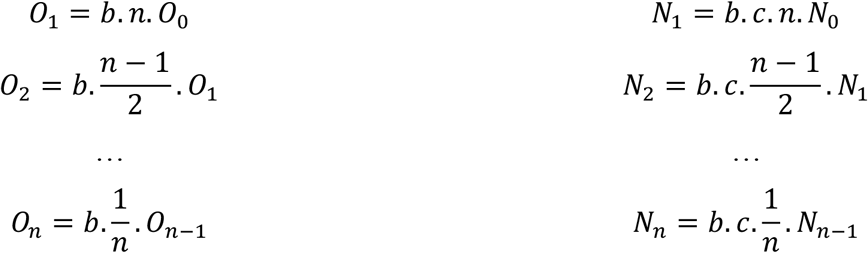

Therefore, *total number of sites in O state* = *n* (*O*_*O*_ + *O*_1_ + ⋯ +*O*_*n*_)

And *no. of R sites bound* = *O*_1_ + 2*O*_2_ + ⋯ + *nO*_*n*_

Similarly, *total number of sites in O state* = *n*(*N*_*O*_ + *N*_1_ + ⋯ + *N*_*n*_)

And *no. of R sites bound* = *N*_1_ + 2*N*_2_ + ⋯ + *nN*_*n*_

Fraction of total (open and nucleosomal) sites bound

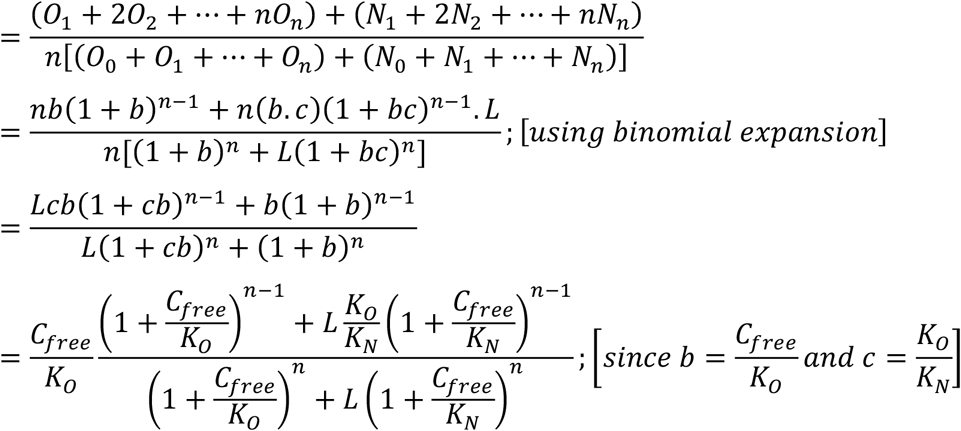

Therefore,

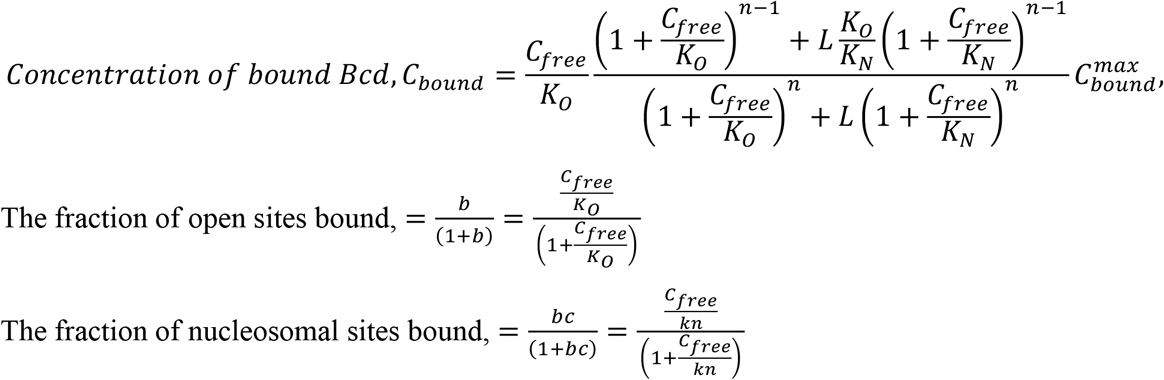

### MWC Model fitting

For the correlated population, we fitted the MWC model explained above to the data.

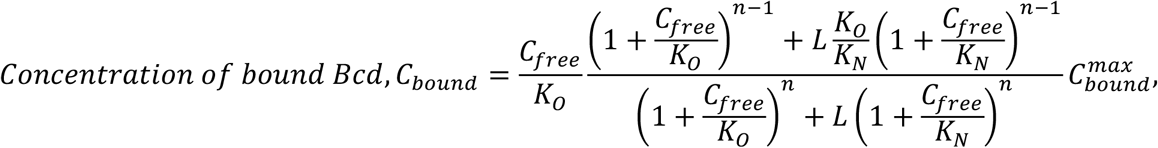

We set the parameters *n, L, K*_*N*_, *K*_*O*_, and 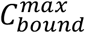 as free and used Brute-force parameter screening within ranges mentioned in Table S1. The capacity 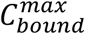 was determined analytically. The threshold for fitted parameters was determined using the nonlinear regression F-test criterion.

To obtain higher resolution, we repeated the screen with finer parameter mesh based on the results of the first screen (*e*.*g*., setting *n* = 2) (Fig. S3B).

### Estimation of capacity

The number of Bcd binding sites that are available for binding determined from the TFBS enriched in the Bcd-bound regions are ~ 2680^23,77^. The nuclear volume is ~ 300 µm^3^ at nc 14^78^. Thereby the concentration of the binding sites for the diploid *Drosophila* embryo at nc 14 is ~ 29.7 nM.

### Simplified MWC model

The MWC model can be reduced to ignore the *K*_*N*_ terms as *K*_*N*_ ≫ *K*_*O*_.

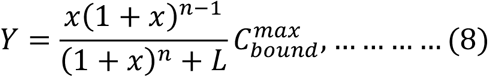

Where, 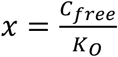 and n can be fixed at 2. Similar parameter screening (mentioned in the *Model fitting* above) was carried for using *n* = 2.

For the comparison of location of the inflection point and the slope at that point, we focus on the parameter sets within threshold for the MWC model and having the upper 10% of *K*_*N*_ values. For each parameter set, we determined the inflection point by calculating slopes at each point of the curve and determining the point with the highest slope for both the MWC model and the simplified model. The values obtained using both the models were compared.

### L versus 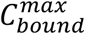 relationship

Step 1: Slope is calculated for all the MWC parameter sets that fit the data within a threshold using the simplified model and the maximum slope is determined for each parameter set.

Step 2: For the simplified model, the slope is given by:

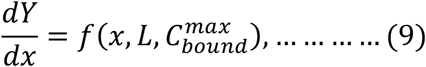

At the inflection point,

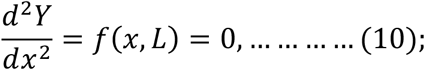

We calculated x corresponding to a range of L values using Eq. 10 and calculated the corresponding 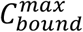 by plugging in the values of x, L, and 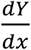 (median of the maximum slopes determined in step 1) in Eq. 9, which ultimately demonstrated the relationship between L and 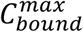.

The fraction of sites bound can be obtained from Eq. 8

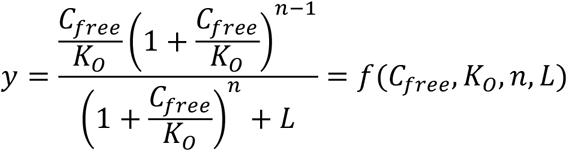

Now,

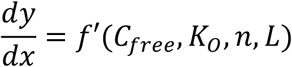

At the inflection point,

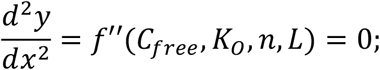

Thus, at highly constrained values of *K*_*O*_ and *n*, the location of the inflection point in the dose/response and the slope at that point are determined by *L*.

### Use of MS2 data

We used the fraction of active nuclei and the transcriptional onset time as reported by Degen et al. ^24^. The RNAP loading rate data is as reported by Eck et al.^49^. MS2 data from references^24,49^ were downloaded as described therein. The fraction of active nuclei as plotted in Figs. 4,5 and S5 were calculated as follows from data published in^24^. First, as described in ^24^, the nc13 MS2 time traces for each nucleus in each embryo were interpolated onto a regular time mesh, noise filters were applied, then each nucleus was placed onto a regular AP coordinate grid according to its average AP coordinate across nc13 interphase. After these steps, the fraction of active nuclei for each embryo at each time point and AP coordinate was calculated as the number of “on” nuclei divided by the total number of nuclei. A nucleus was counted as “on” if its filtered MS2 intensity was greater than zero. The mean fraction of active nuclei as a function of AP coordinate (plotted in Figs. 4,5 and S5) were averaged across all embryos and from time points t = 420 to 820, the time window during which the fraction of active nuclei appeared to stabilize.

The onset time was used directly from the data downloaded as described in ^24^ (see Fig. S1 within that reference).

The RNAP loading rate was used directly from the published graphs in ^49^.

### Thermodynamic models of transcription

Thermodynamic models are based on the idea that the gene expression level can be deduced from the probabilities that species such as activators, repressors, RNAP are bound to the regulatory region associated with the gene of interest^53,54^. The probability of regulatory region occupancy by a species (TF and RNAP in our case) is given by the ratio of sum of weights of favorable microstates and that of weights of all the microstates^57^. The weights can be written in terms of the TF concentrations and the dissociation constant^49,57^.

For easy visualization, we consider a simple case in which two Bcd binding sites are occluded by a nucleosome in the inactive state (Fig. 4A). In our model, Bcd can bind in the open as well as in the nucleosomal state of the DNA with dissociation constant *K*_*O*_ and *K*_*N*_ where *K*_*O*_ ≪ *K*_*N*_. The unbound open and nucleosomal states are in equilibrium, L being the equilibrium constant. However, transcription occurs only when the DNA is in the open state and Bcd is bound. Therefore, the probability of finding an open state.

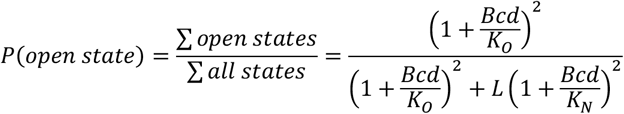

In case of the two nucleosomes in *hbP2* which occlude clusters of 5 and 4 sites, binding of Bcd to each cluster are independent events and the probability of finding open state is therefore product of the probabilities finding both the clusters in open state.

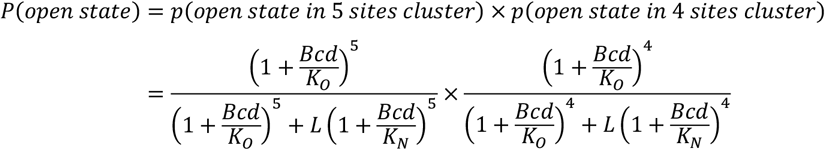

The equation can be modified to obtain the probability of transcription based on the assumption of which state is competent for transcription. We have considered three cases as mentioned in Eq. (2-4).

The parameter sets from the fit to the DNA-binding dose/response map are used to simulate the *P*_*active*_ for the three cases. In order to test whether parameter combinations other than the ones from the DNA-binding dose/response map could result in the *hb* pattern, we fitted the models (Eq. 2-4) using the least-squares fitting algorithm lsqcurvefit and the global optimization solver MultiStart in Matlab.

For the RNAP loading rate, RNAP binding to the promoter needs to be taken into account, in addition to the previously mentioned microstates (Fig. 5D). In the simplified case of two Bcd binding sites occluded by a nucleosome, the probability of RNAP binding is given by:

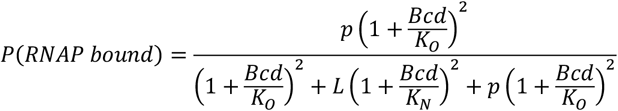

Where, *p* = *RNAP*/*K*_*p*_.

Similarly, considering **Case 1** for *hbP2* it can be extended to

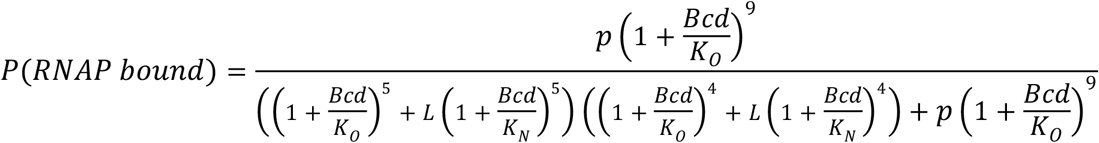

### Promoter state model

Here we consider the chemical master equation formalism to calculate the probability distribution of promoter states. We define *P*_*i*_ as the probability that the promoter is in State *i*. The transition to State *i* can occur in two ways:

1. Being in State *i* − 1 and transitioning “to the right” (towards the “on” State) with rate constant *k*_*on*_(*Bcd*), or
2. Being in State *i* + 1 and transitioning “to the left” (towards the “fully off” State) with rate constant *k*_*off*_

Here, *k*_*on*_(*Bcd*) = *k*_*on*,0_*P*_*active*_. The formula for *P*_*active*_ varies based on the assumption of which Bcd bound state can drive activation, and we consider the three cases mentioned in Eq. (2-4).

Similarly, transitioning out of State *i* can occur in two ways:

1. Transitioning “to the right” with rate constant *k*_*on*_(*Bcd*), or
2. Transitioning “to the left” with rate constant *k*_*off*_

For the generalized case of a promoter with N states (N-1 OFF states and one ON state), for the States not equal to State 1 (the “fully off” state) or State N (the “on” state):

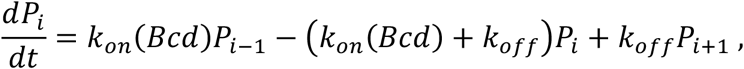

If i = 1:

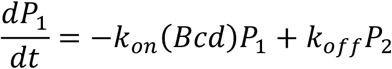

If i = N:

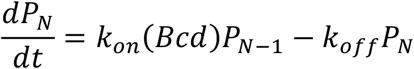

The vector formulation is:

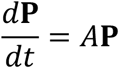

Here, **P** is a vector of the probabilities of occupying the respective N states, and *A* is the matrix of coefficients:

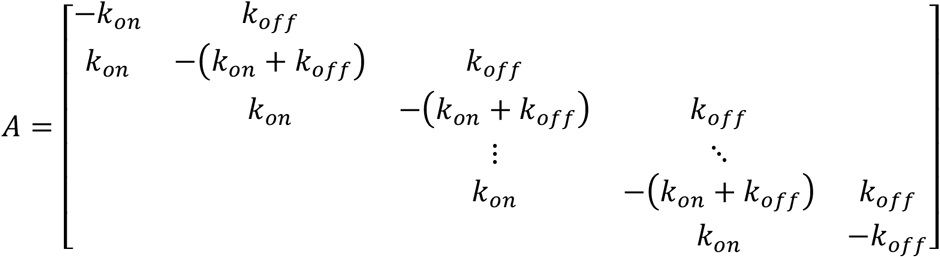

…where, for simplicity, we have removed the explicit Bcd dependence of *k*_*on*_.

These equations can be non-dimensionalized by dividing by *k*_*off*_:

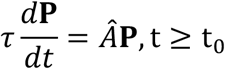

…where *τ* = 1/*k*_*off*_ is the system time scale and t_0_ is the mitotic repression window. The normalized matrix Â is:

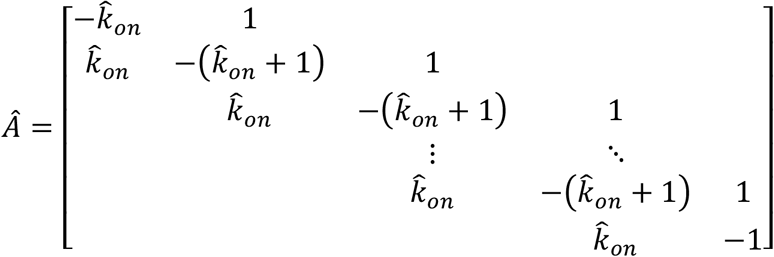

… where 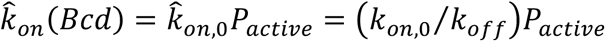.

The initial condition is:

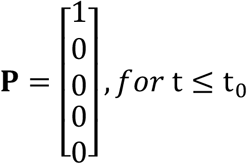

At steady state, these equations reduce to:

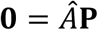

Where the LHS is the zero vector of ℝ^*N*^ and **P** is a vector of the probabilities of occupying the respective N states. In this formulation, the equations are linearly dependent. Noting that the probabilities of all states sum to 1, to find the steady state, we can therefore replace the final equation by a row of 1’s, and the final element of the vector on the RHS with a 1 (i.e, the sum of the probabilities of all states).

The dimensionless parameter 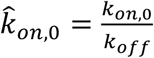 can be determined by fitting this model at steady state to the fraction of active nuclei ^24^. The two additional adjustable parameters (the system time scale, *τ*, and the mitotic repression window, *t*_0_) can be determined by fitting the non-steady state model to the onset times. To do this, we defined the onset time as the time it takes for the nuclei at to reach 50% of the steady state value.

## Author contributions

Conceptualization: S.S.D. and G.T.R. Imaging: S.S.D. Analysis: S.S.D. and G.T.R. Investigation: S.S.D. and G.T.R. Visualization: S.S.D. and G.T.R. Supervision: G.T.R. Writing-original draft: S.S.D. and G.T.R.

