## Supplemental Figures (S1-S5), Supplemental table S1 for "Genome-wide mapping of Bicoid/DNA interactions reveals quantitative constraints on transcriptional regulation"

### Supplementary materials

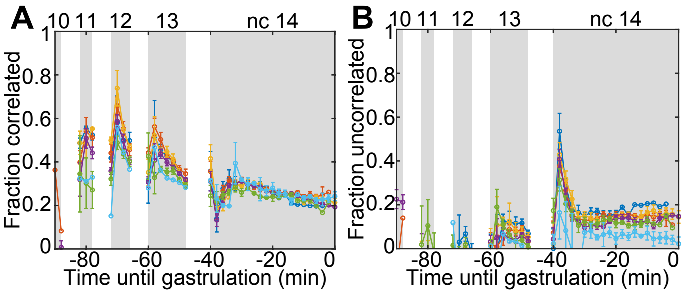


**Figure S1:** Dynamics of the fraction of immobile Bcd populations at different locations with each color denoting a particular location of the ROI (following the same scheme as in Fig. 1B). (A) Dynamics of the fraction of population correlated with His2Av-RFP. (B) Dynamics of the fraction of population uncorrelated with His2Av-RFP. Data points, mean; error bars, SEM.

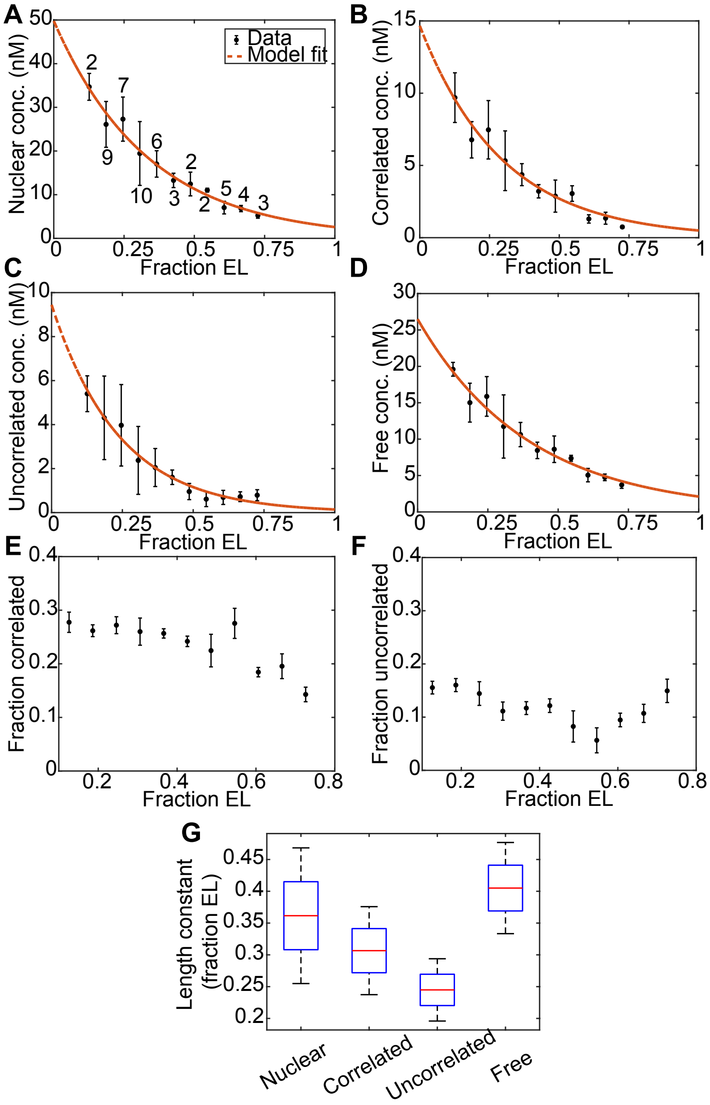


**Figure S2:** Spatial variation of the populations of Bcd during nc 14 determined using tRICS. (A-D) Exponential decay fit to the AP profile of each population including total nuclear concentration (A), concentration of the correlated population (B), concentration of the uncorrelated population (C), concentration of the free population (D). Data points, mean; error bars, SEM. Numbers on the curve in the (B) indicate number of embryos imaged for each position. (E-F) Spatial variation of the fraction of immobile populations including fraction of correlated population (E) and fraction of uncorrelated population (F). (G) Comparison of the length constant for the exponential decay fit to each population.

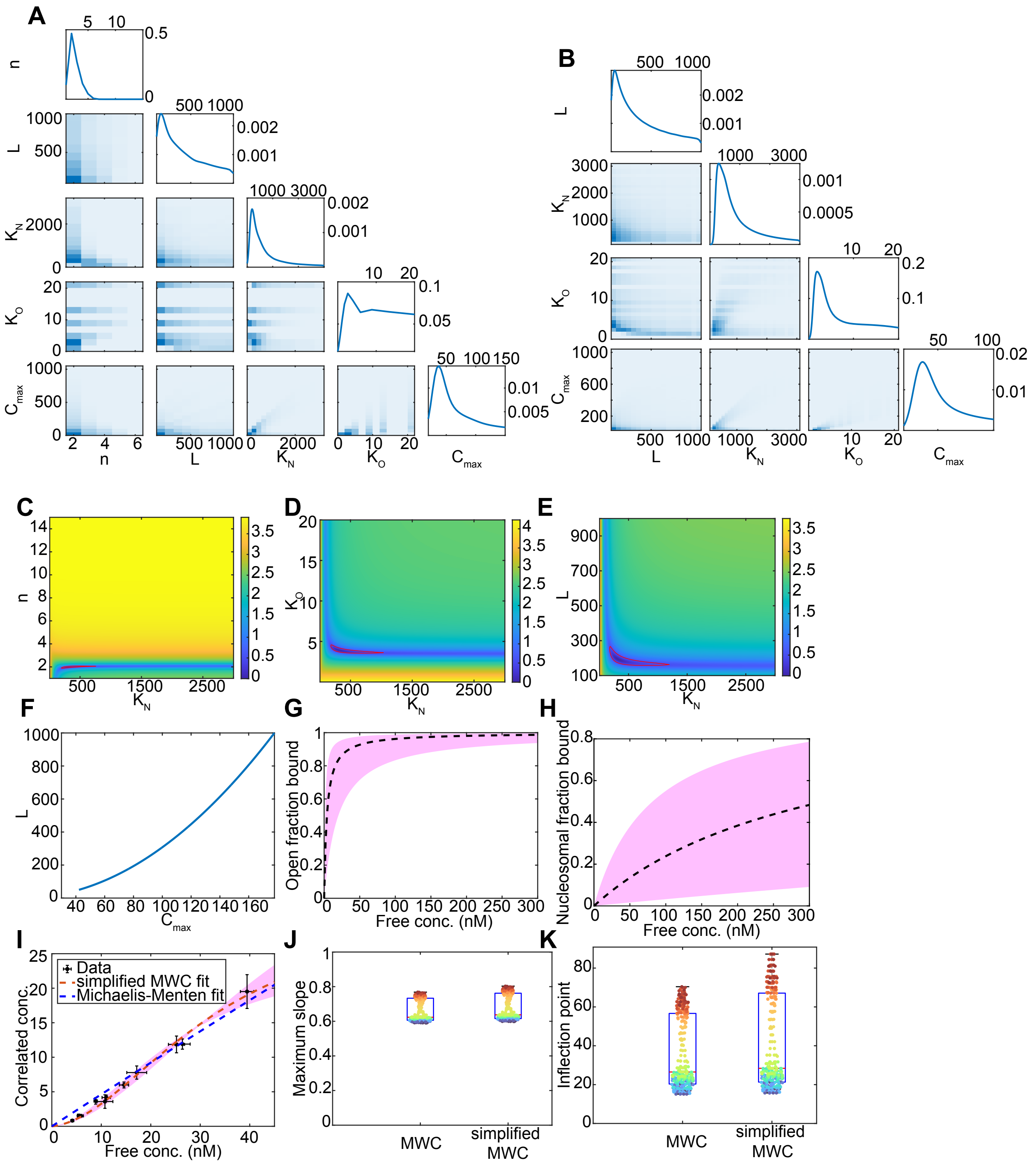

**Figure S3:** MWC model fit to the dose/response map between free and correlated population using brute force parameter screening. (A-B) Corner plot showing the distribution of parameters that fit the dose response map (pink region in Fig. 3B) with initial screening varying all the parameters (A) and screening using a finer mesh and fixed $n = 2$ based on the results of the first screen (B).

(C-E) Heatmaps showing the correlation between parameters, for each plot the values of the parameters other than the variables are fixed at the those in the parameter set that gives the lowest SSE ($n$ = 2, $L = 194$, $K_N = 320$, $K_O = 4$, $C_{max} = 52$), correlation between $n$ and $K_N$ (C), correlation between $K_o$ and $K_N$ (D), correlation between $L$ and $K_N$ (E). Colorbars, log normalized SSE relative to the global minimum. (F) Correlation between $L$ and $C_{max}$, regardless of the changes in other parameters. (G-H) Fraction of sites bound. Dashed line indicating the curve generated using the parameter set that gives the lowest SSE and the shaded region indicating all the parameter sets (pink region in Fig. 3B) including fraction of open sites bound (G) and fraction of nucleosomal sites bound (H). (I) Michaelis-Menten and simplified MWC fit to the dose/response between free Bcd and correlated Bcd. The pink shaded region indicates the MWC fit using different parameter sets that fit the data within a set threshold. Data points, mean; error bars, SEM. (J-K) Comparison between the MWC model and the simplified MWC model for maximum slope (J) and location of the inflection point (K). Same color of points represents the values calculated for the same parameter sets in both the models.

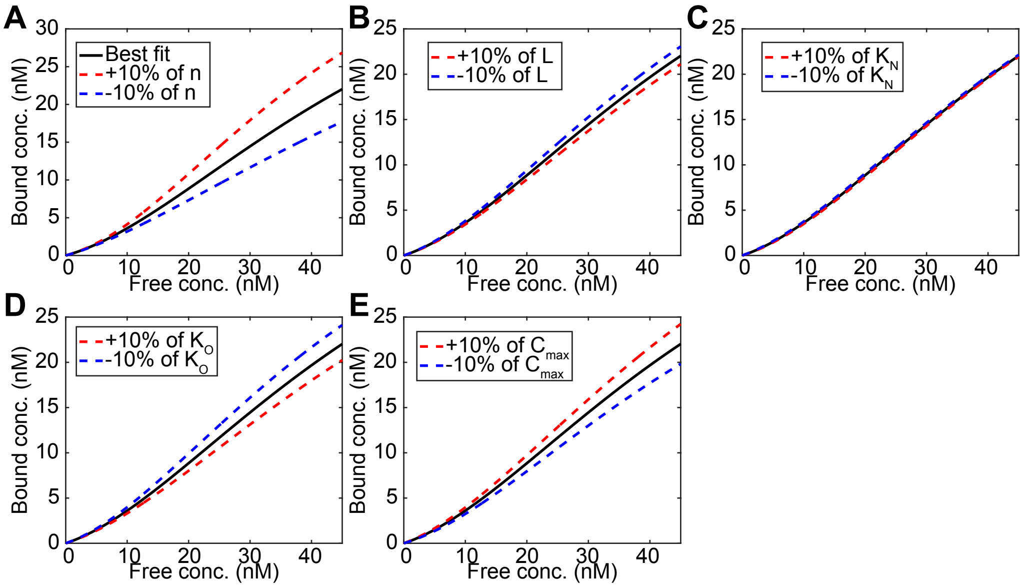


**Figure S4:** (A-E) Sensitivity analysis to show the change in dose/response resulting from 10% changes in the parameters $n$ (A), $L$ (B), $K_N$ (C), $K_O$ (D), $C_{max}$ (E), while keeping all other parameters constant. Solid black line represents the prediction corresponding to the parameter set ($n$ = 2, $L = 194$, $K_N = 320$, $K_O = 4$, $C_{max} = 52$).

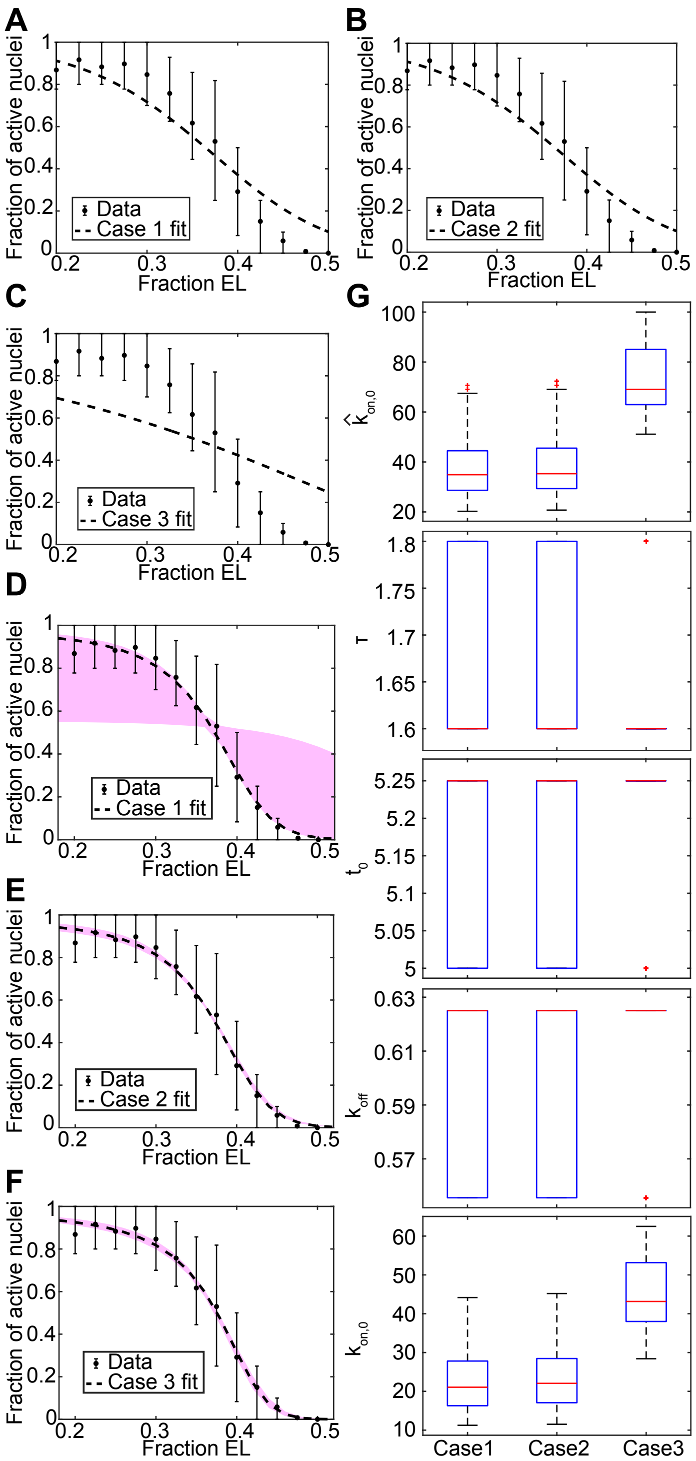


**Figure S5:** Transcriptional dynamics driven by *hbP2*. (A-C) Enhancer occupancy by Bcd model fit to the fraction of active nuclei using nonlinear least-squares fitting for Case 1 (A), Case 2 (B), Case 3 (C). Dashed lines represent the best fit. (D) Model fit to the fraction of active nuclei using

Case 1. Dashed line represents the best fit and the pink region represents the fits using all the parameter sets in Fig. 3B. (E-F) Reversible promoter state model fit to the fraction of active nuclei using Case 2 (E) and Case 3 (F). Dashed line represents the best fit and the pink region represents the fits using parameter set in Fig. 3B that fit the fraction of active nuclei within a threshold. In (A-F), data points, mean; error bars, inner 68th percentile range. (G) Comparison of parameter values determined by fitting the three cases of the reversible promoter state model to the fraction of active nuclei and transcriptional onset time.

**Table S1**

| Parameter | Ranges | Source |
|---|---|---|
| $n$ | 1-15 | Upper limit based on number of Bcd binding sites in enhancers[7] |
| $L$ | 100-1000 | [44] |
| $K_N$ | 6-3000 | [44] (based on the ratio between $K_O$ and $K_N$) |
| $K_O$ | 0.6-20 | [11,24,45,46] |
